# Chromosome-scale genome assembly of the pink bollworm, *Pectinophora gossypiella*, a global pest of cotton

**DOI:** 10.1101/2022.10.07.511331

**Authors:** Amanda R. Stahlke, Jennifer Chang, Sivanandan Chudalayandi, Chan C. Heu, Scott M. Geib, Brian E. Scheffler, Anna K. Childers, Jeffrey A. Fabrick

## Abstract

The pink bollworm, *Pectinophora gossypiella* (Saunders) (Lepidoptera: Gelechiidae), is a major global pest of cotton. Current management practices include chemical insecticides, cultural strategies, sterile insect releases, and transgenic cotton producing crystalline (Cry) protein toxins of the bacterium *Bacillus thuringiensis* (Bt). These strategies have contributed to eradication of *P. gossypiella* from the cotton growing areas of the United States and northern Mexico. However, this pest has evolved resistance to Bt cotton in Asia, where it remains a critical pest, and the benefits of using transgenic Bt crops have been lost. A complete annotated reference genome is needed to improve global Bt resistance management of the pink bollworm. We generated the first chromosome-level genome assembly for pink bollworm from a Bt-susceptible laboratory strain (APHIS-S) using PacBio continuous long reads for contig generation, Illumina Hi-C for scaffolding, and Illumina whole-genome re-sequencing for error-correction. The psuedohaploid assembly consists of 29 autosomes and the Z sex chromosome. The assembly exceeds the minimum Earth BioGenome Project quality standards, has a low error-rate, is highly contiguous at both the contig and scaffold level (L/N50 of 18/8.26 MB and 14/16.44 MB, respectively), and complete, with 98.6% of lepidopteran single-copy orthologs represented without duplication. The genome was annotated with 50% repeat content and 14,107 protein-coding genes, further assigned to 41,666 functional annotations. This assembly represents the first publicly available complete annotated genome of pink bollworm and will serve as the foundation for advancing molecular genetics of this important pest species.

## Introduction

The pink bollworm *Pectinophora gossypiella* is globally one of the most damaging insect pests of cotton (CABI, 2022; Henneberry & Naranjo, 1998). Transgenic cotton producing *Bacillus thuringiensis* (Bt) protein toxins is used by many countries to target and kill insect pests including the pink bollworm, thereby providing both economic and environmental benefits (ISAAA, 2019; National Academies of Sciences & Medicine, 2016; Sanahuja et al., 2011). However, the evolution of pest resistance threatens the continued success of such Bt crops.

The pink bollworm was successfully eradicated from the cotton growing areas of the U.S. and northern Mexico (Perdue, 2018), in large part because it remained susceptible to Bt transgenic cotton that simultaneously produced two different Bt crystalline (Cry) toxins (Tabashnik et al., 2021). Pink bollworm in China have also remained susceptible to Bt cotton producing Cry1Ac, due primarily by the mass planting of F_2_ hybrid cotton that provided sufficient non-Bt refuge plants that countered the evolution of resistance (Wan et al., 2017). However, pink bollworm have evolved practical resistance to Bt cotton producing Cry1Ac in Pakistan (Akhtar et al. 2016; Akhtar et al. 2020) and both Cry1Ac and Cry2Ab in India (Dhurua & Gujar, 2011; Naik et al., 2018).

To date, pink bollworm resistance to the Bt toxins Cry1Ac and Cry2Ab has been associated with the midgut cadherin *PgCad1* and the ATP-binding cassette *PgABCA2* genes, respectively. For Cry1Ac, a total of 20 mutations in *PgCad1* have been identified in resistant pink bollworm from China, India, and the U.S. (Fabrick et al., 2014; Fabrick & Tabashnik, 2012; Morin et al., 2003; Wang et al., 2019a; Wang et al., 2018; Wang et al., 2022). Resistance to Cry1Ac in a laboratory strain of pink bollworm from Arizona was also found to involve reduced transcription of the *PgCad1* gene leading to markedly lower abundance of the PgCad1 Cry1Ac receptor protein (Fabrick et al., 2020b). Resistance to Cry2Ab has been shown to involve mutations in the ATP-binding cassette *PgABCA2* gene in pink bollworm laboratory strains from Arizona and field populations from India (Mathew et al., 2018). Genetic knockout of *PgABCA2* by CRISPR/Cas9 gene editing in a susceptible lab strain (APHIS-S) causes resistance to Cry2Ab (Fabrick et al., 2021). In addition to mutations in *PgABCA2* imparting resistance to Cry2Ab, a second unknown mechanism of resistance also appears be present in one of the Arizona strains (Fabrick et al., 2020a). Having a complete and fully annotated genome assembly facilitate further identifying and physically mapping additional Bt resistance loci, and may also uncover novel targets for future pest management strategies.

Here, we present the first high-quality, chromosome-scale, annotated genome assembly of pink bollworm from a Bt-susceptible laboratory strain (APHIS-S) using the best practices for assembly and error-correction of Pacific Biosciences (PacBio) continuous long reads (CLR), recently implemented for insect pests (Chang et al., 2022; Rhie et al., 2021; Stahlke et al., 2022).

## Methods

The APHIS-S strain of pink bollworm has been maintained under laboratory conditions for more than 40 years without exposure to Bt toxins or other insecticides (Bartlett, 1995; Liu et al., 2001). APHIS-S was derived from the APHIS wild type strain maintained by USDA APHIS Center for Plant Health Science and Technology (CPHST) (Phoenix, AZ), and was used for field releases as part of the Sterile-Insect Technique program in the San Joaquin Valley of California and the Binational Pink Bollworm Eradication Program (Grefenstette et al., 2009). Insects were received from USDA APHIS CPHST in 2017 and larvae were reared at USDA ARS (Maricopa, AZ) on wheat germ diet (Bartlett & Wolf, 1985). All insect life stages were maintained at 26°C, 14 h light:10 h dark.

DNA extraction was performed from a single female 4^th^ instar larva (ToLID ilPecGoss1) using the NucleoMag Tissue kit (Macherey-Nagel, Allentown, PA, USA). The high molecular weight DNA was prepared for PacBio sequencing using the SMRTbell Express Template Prep Kit 2.0 for Continuous Long Read (CLR) generation (Pacific Biosciences, Menlo Park, CA, USA). Sequencing was performed on a Sequel II System (Pacific Biosciences) using Binding Kit v2.0, Sequencing kit v2.0, and SMRT Cell 8M. To target CLR reads, the library was sequenced using a 20-hour movie time on one SMRTcell. Short-read sequencing data for error correction was generated from the same individual using the TruSeq Nano DNA Low Throughput Library Prep Kit and IDT for Illumina – TruSeq DNA UD Indexes (Illumina, San Diego, CA, USA). Standard protocols were followed to yield a paired library with a 350 bp insert size and 50 bp standard deviation. Paired end 150 bp sequence reads were generated on a NovaSeq 6000 System (Illumina) at the HudsonAlpha Genome Sequencing Center (Huntsville, AL, USA).

DNA from a second female 4^th^ instar larva (ToLID ilPecGoss2) was crosslinked using the Arima Hi-C low input protocol, and proximity ligation was performed using the Arima Hi-C Kit (Arima Genomics, Carlsbad, CA, USA). After proximity ligation, the DNA was sheared using a Diagenode Bioruptor and then size-selected to enrich for DNA fragments of 200-600 bp. An Illumina library was prepared from the sheared and size-selected DNA using the Accel-NGS 2S Plus DNA Library kit (Swift Biosciences, Ann Arbor, MI, USA). The final Illumina Hi-C library was sequenced on a NovaSeq 6000 System at the HudsonAlpha Genome Sequencing Center (Huntsville, AL, USA).

The primary pseudohaploid and corresponding alternate genome assemblies were generated following established best practices, implemented as the NextFlow pipeline, polishCLR (Chang et al., 2022; Stahlke et al., 2022). Briefly, contigs were generated from the raw FASTQ PacBio CLR subreads with the FALCON assembler and FALCON-Unzip (Chin et al., 2016) using the pb-assembly conda environment v.0.0.8.1, which includes one-round of haplotype-aware polishing by Arrow (Pacific Biosciences). Duplicate haplotypes were removed from primary and alternate contigs with purge_dups v1.2.5 (Guan et al., 2020), using coverage cutoffs automatically calculated from histograms of PacBio coverage.

Paired-end Hi-C reads were aligned to the purged primary assembly with bwa-mem v2.2.1 (Li, 2013), allowing for chimeric alignments with the −5SP options. Alignments were converted to a contact-map visualization for manual curation in Juicebox v1.111.08 (Durand et al., 2016), where contigs were joined, broken, and re-oriented, and duplicate haplotypes were removed. The scaffolded primary, purged alternate, and mitochondrial genome (see below) were then provided as input to step two of the polishCLR pipeline for an additional round of polishing with Arrow using the PacBio reads and consensus variant recalling with Merfin v1.0 (Formenti et al., 2021a), followed by two rounds of error-correction with FreeBayes v1.0.2 (Garrison & Marth, 2012) using the re-sequencing Illumina reads generated from ilPecGoss1. Scaffolded and polished primary and alternate scaffolds were screened for cobionts by visualizing coverage, GC-content, and BLAST (Altschul et al., 1997) and DIAMOND (Buchfink et al., 2015) hits with BlobTools (Challis et al., 2020).

We assembled, polished, and automatically annotated the mitochondrial genome using Ag100MitoPolishCLR (https://github.com/Ag100Pest/Ag100MitoPolishCLR) (Allio et al., 2020; Formenti et al., 2021b), as previously described for *H. zea* (Stahlke et al., 2022). We seeded the assembly with the mitochondrial genome of a closely-related moth of the same tribe, Pexcopiini, *Sitotroga cerealella* (Lepidoptera: Gelechiidae: Apatetrinae) (Yuan et al., 2019, NC_041123.1). Mitochondrial annotations were manually reviewed (Cameron, 2014) and corrected in Geneious Prime v2021.1.1 (Kearse et al., 2012).

We surveyed the primary assembly for sex chromosomes, using the chromosome quotient method (Hall et al., 2013) to identify candidate W-linked scaffolds, and synteny to validate the Z chromosome. The chromosome quotient method employs the assumption that sex chromosomes in a ZZ/ZW system will have half the coverage with heterogametic sample re-sequencing. We aligned forward reads of the Illumina polishing data back to the scaffolds with bwa mem, then used the idxstats function in SAMtools v1.9 (Li et al., 2009) to calculate average coverage across each scaffold. We then identified the mean coverage, normalized by length of the first five scaffolds, and identified scaffolds that showed half coverage relative to the mean (Hall et al., 2013). To validate the assignment of the Z chromosome, we visualized synteny between the genome of the most closely related species with a chromosome-scale assembly with an annotated Z chromosome, *Agonopterix subpropinquella* (Lepidoperta: Depressariidae) (GCA_922987775.1) and ilPecGoss1. At the time of writing, there were not any W scaffolds or markers publicly available within the Gelechiidae family.

GenomeScope 2.0 (Ranallo-Benavidez et al, 2020) was used to estimate genome size using a 21-mer k-mer histogram of the Illumina re-sequencing reads constructed with KMC v. 3.1.0, allowing a maximal coverage of 100 million k-mers. Additional valuations of the assembly were performed with KAT v2.4.1 (Mapleson et al, 2017) to assess assembly composition and quality, BUSCO 5.2.2 (Manni et al., 2021) to assess completeness, and Merqury to assess the base accuracy of the assembly.

Autosomal chromosome-scale scaffolds of the final primary assembly were ordered according to size, followed by the Z chromosome, and unassigned contigs according to size. Repetitive elements were identified using RepeatMasker v4.1.0 (http://repeatmasker.org) with a *de novo* library of repeats built with RepeatModeler v2.0.2 (Flynn et al., 2020). The final primary assembly was submitted to the National Center for Biotechnology Information (NCBI) for automated eukaryotic genome annotation (Thibaud-Nissen et al. 2016), where reference sequence (RefSeq) gene models were constructed both *ab initio* and from all publicly available RNAseq evidence of pink bollworm (https://www.ncbi.nlm.nih.gov/genome/annotation_euk/process/) (PRJNA450266; PRJNA528157; Tassone et al., 2016). From these gene models, functional associations to gene ontologies and pathways were assigned using the AgBase Functional Annotation Workflow (Saha et al., 2021).

## Results & Discussion

From the first female larva, ilPecGoss1, we generated 112.7 GB of long-read PacBio CLR data (ToLID ilPecGoss1) and 143.57 million read pairs of Illumina 150 bp re-sequencing data (43 Gb). From the second female larva, ilPecGoss2, we generated 10.7 million read pairs of 150 bp Hi-C data.

We produced and annotated a complete mitochondrial genome for ilPecGoss1, complementary to a previously released assembly (KM225795; Zhao et al., 2016). At 15,209 bp in length, it is seven base pairs longer than the first assembly, but otherwise similar to both KM225795 and the mitochondrial genomes of other Gelechiidae (Timmermans et al., 2014). As is typical among Lepidoptera, one additional copy of both tRNA-Leu and tRNA-Ser were found. The mitochondrial genome was circularized according to the conventional Lepidopteran tRNA organization, with an origin of replication behind the tRNAs MIG (Cameron, 2014; Timmermans et al., 2014).

The only publicly available genome assembly of the pink bollworm prior to our work (JALBXW000000000.1) was constructed entirely of Illumina short-read shotgun data, thereby having low contiguity (scaffold L/N50 of 79,304/1.32 Kb), low completeness (51.2% single, complete orthologs of Lepidoptera odb10), and was not annotated, limiting its utility. Contig assembly of our CLR data yielded a primary pseudohaploid genome 566.98 M in total size, composed of 474 contigs, with a conitg L/N50 of 38/3.70 Mb and a high level of completeness as assessed using the BUSCO Lepidoptera odb 10 ortholog set (C:98.7% [S:89.4%, D:9.3%], F:0.3%, M:1.0%) (Supp. Table S1). After removing duplicate haplotypes, we retained 304 contigs in the primary assembly, improving the L/N50 to 29/5.13 Mb, and reducing duplicate BUSCOs. However, this removal did cause a small decrease in complete BUSCOs (C:98.5% [S:97.8%, D:0.7%], F:0.4%, M:1.1%) (Supp. Table S1).

After manual curation of the scaffolded assembly (Fig. 1A) and completion of the polishCLR pipeline, some gaps were closed, further improving the contig N50 from 4.21 to 8.26 Mb, and the QV from 39.67 to 42.98 (Supp. Table S2). Visualization of BLAST hits on each scaffold of this assembly supported the removal of 31 scaffolds from the primary assembly and one from the alternate assembly, all showing high identity with *Pseudomonas* spp. The final polished primary and alternate assemblies were 480.69 Mb and 322.03 Mb in total length, with L/N50s of and 282/0.36 Mb, respectively (Figure 1C, Table 1). Total genome size estimated from k-mer analysis of re-sequencing reads was about 40 Mb smaller, at 447 Mb (Fig. 1C), but BUSCO (Table 1) and KAT results support our assembly as both complete and nonredundant (Figure 1B).

**Table 1.** Pink bollworm (ilPecGoss1.1) genome assembly and annotation metrics.

| Metric | Value |
| --- | --- |
| **Alternate contig assembly** |  |
| Total number | 1,695 |
| Assembly size (bp) | 322,000,363 |
| L50/N50 (bp) | 282/358,484 |
| L90/N90 (bp) | 942/84,990 |
| Largest contig (bp) | 1,259,928 |
| **Primary assembly** |  |
| *Scaffolds* |  |
| Total number | 151 |
| Size (bp) | 476,494,545 |
| L50/N50 (bp) | 13/16,702,263 |
| L90/N90 (bp) | 28/6,805,748 |
| Largest (bp) | 28,314,428 |
| *Contigs* |  |
| Total number | 242 |
| Size (bp) | 476,485,531 |
| L50/N50 (bp) | 18/8,257,054 |
| L90/N90 (bp) | 73/12,44,681 |
| Largest (bp) | 17,914,429 |
| **Lepidoptera odb10 BUSCOs** |  |
| Complete (C) | 5,539 (98.5%) |
| Complete single copy (S) | 5169 (97.8%) |
| Complete duplicated (D) | 370 (0.7%) |
| Fragmented (F) | 211 (0.4%) |
| Missing (M) | 58 (1.1%) |
| **Gene structural annotations** |  |
| Genes (total) | 16,460 |
| Protein coding | 14,107 |
| Non-coding | 2,252 |
| Pseudogenes | 101 |
| RefSeq mRNAs | 23,439 |
| RefSeq proteins | 23,452 |
| Repeat content bp (%) | 52.98% |

**Figure 1.**
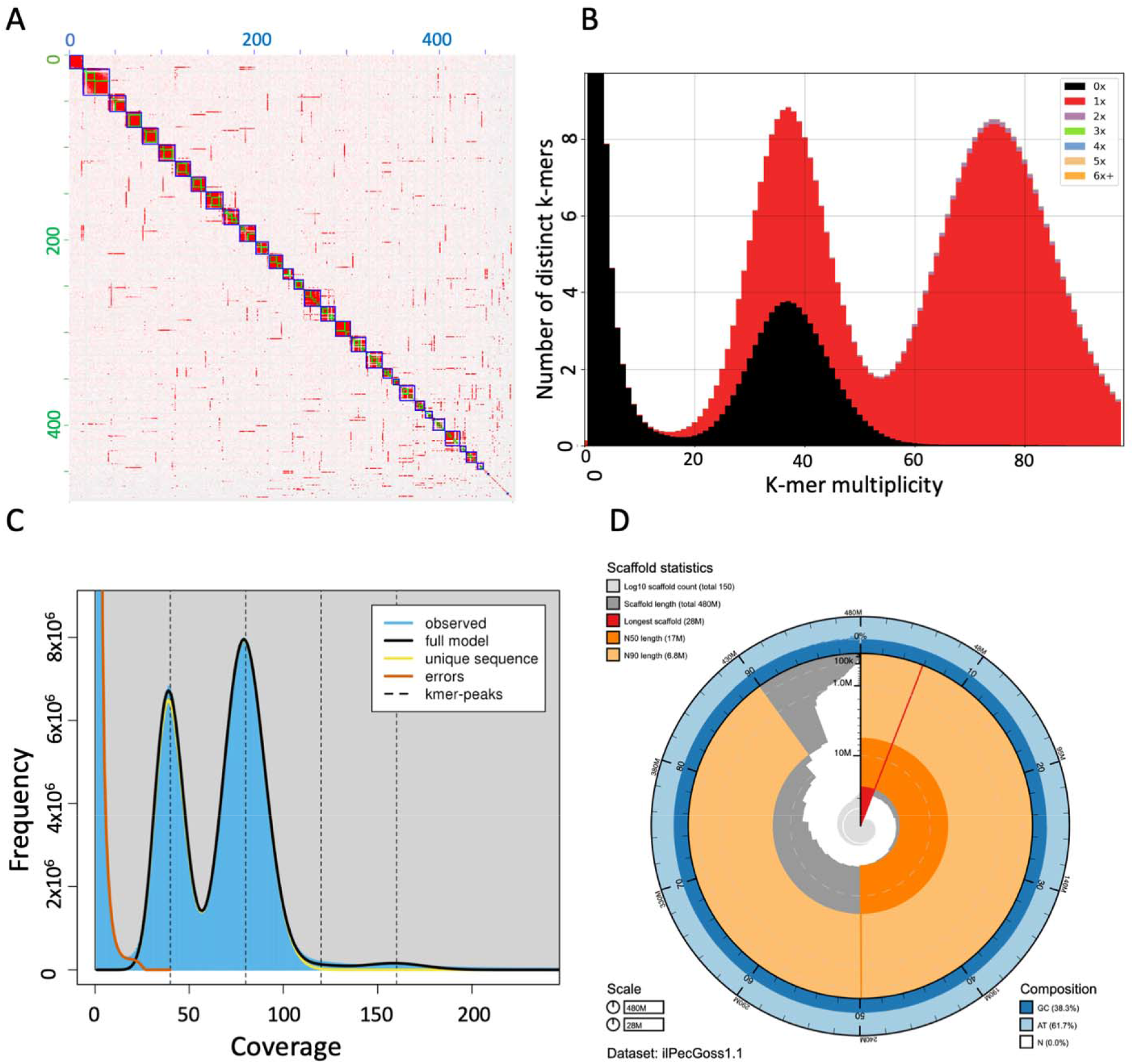
Features of the pink bollworm genome assembly. *(A)* Hi-C contact map adjoining and ordering 30 pseudochromosomes after the manual curation step. *(B)* K-mer comparison of Illumina resequencing reads versus the final primary assembly. *(C)* K-mer visualization of Illumina resequencing reads to estimate genome size, heterozygosity, and repetitive content. *(D)* Snailplot indicating metrics of the primary assembly including scaffold N50, N90, and GC content.

The Z chromosome was apparent upon viewing the Hi-C contact map in Juicebox as a chromosome-scale scaffold with approximately half the coverage of contact mappings relative to the 29 chromosome-scale autosomes (Fig. 1A). We validated the identity of the Z chromosome as the scaffold sharing high identity with the Z chromosome of *A. subpropinquella* (Fig. 2) as well as being the largest chromosome-scale scaffold with roughly half coverage (Supp. Fig. S1) compared to the other chromosomes. 74 smaller contigs also had approximately half coverage, indicating they could be fragments of the Z or W chromosomes (Supp. Table S3, Fig. S1). Future analyses should consider these scaffolds as potentially sex-linked.

**Figure 2.**
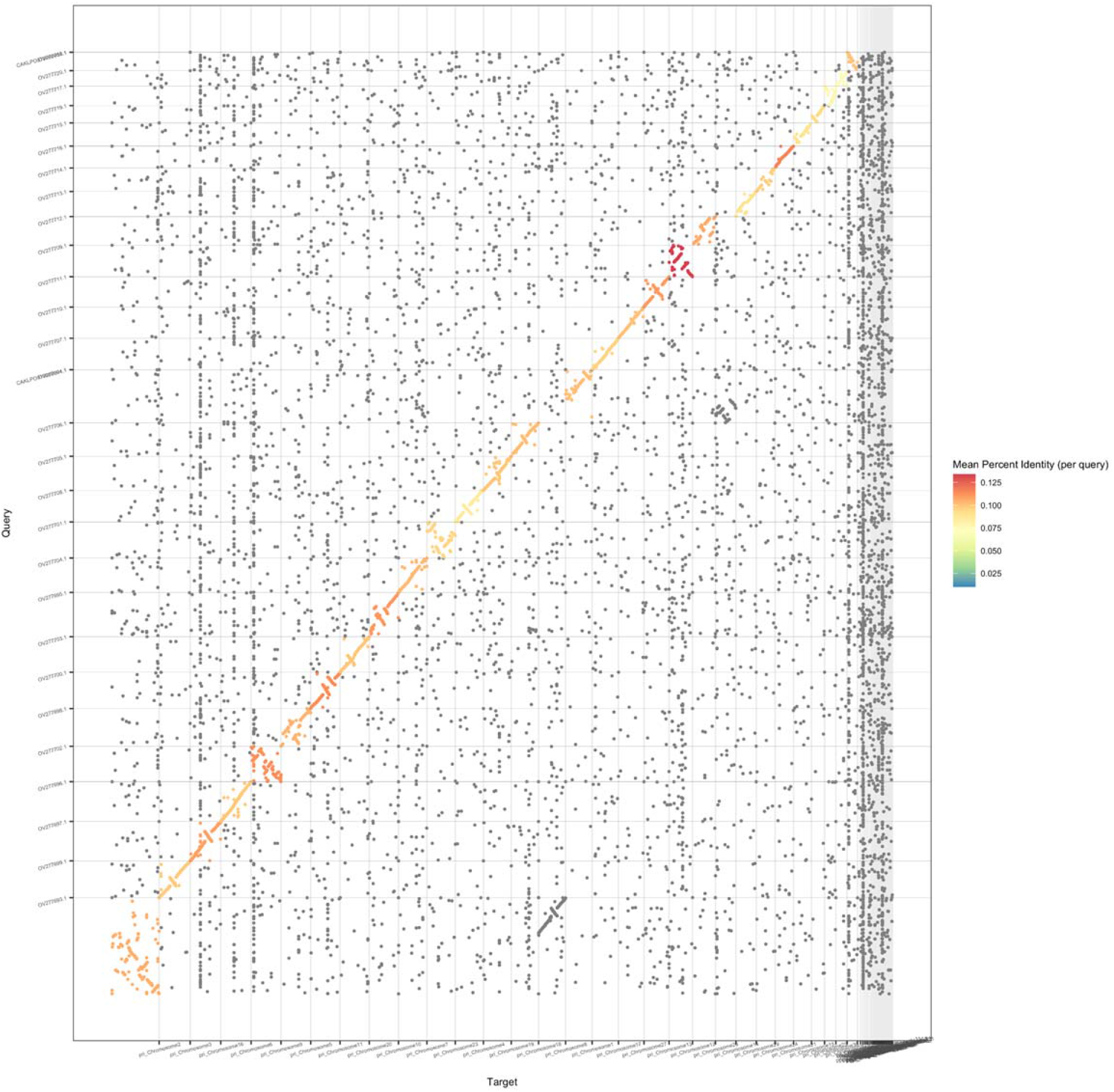
Synteny dot plot between the query genome, *Agonopterix subpropinquella* (GCA_922987775.1), and our submitted pink bollworm genome (ilPecGoss1.1) as the target, with the Z chromosome listed first in both assemblies.

Repeatmasker identified approximately 53% of the genome as repetitive elements, with the plurality (23.83% of the genome) classified as retrotransposons (Supp. Table S4). This is relevant as transposon insertions in the cadherin gene *PgCad1* are linked with resistance to Cry1Ac in pink bollworm from both the U.S. and China (Fabrick et al., 2011; Li et al., 2019; Wang et al., 2019b). Cry1Ac-resistant pink bollworm from India also harbor insertions in *PgCad1* that share sequence similarity with several transposable elements (Fabrick et al., 2014), but have not been fully characterized. These data suggest that mobilization of extant transposable elements plays an important role in the evolution of resistance to Bt crops in pink bollworm.

The NCBI Eukaryotic Genomic Annotation Pipeline predicted 16,460 genes, of which 14,107 protein coding genes gave rise to 23,439 RefSeq transcripts (Table 1). Approximately 90.1% of RefSeq models were supported by RNA-seq evidence. Of the protein coding genes, 13,803 proteins (97.8%) were assigned functional annotations associated with gene ontologies or InterPro, and 8,843 (62.7%) were assigned to a KEGG pathway. These gene ontology and pathway assignments may prove especially useful in dissecting mechanisms of Bt resistance because polygenic adaptation is predicted to be likely in cases of evolution from standing variation (Gould, 1998; Taylor et al., 2021).

Our assembly of *P. gossypiella*, ilPecGoss1.1, exceeds the Earth BioGenome Project minimum reference standard of 6.C.Q40 (Lawniczak et al., 2022; Lewin et al., 2018). This assembly and the associated annotations will provide a critical foundation to advance resistance genomics. The molecular mechanisms of resistance to Bt Cry toxins in pink bollworm are known to range from relatively simple mutations in Bt receptors genes, to complex regulation of gene expression and/or mRNA splicing, and altered protein trafficking (Fabrick et al., 2020b; Fabrick et al., 2014; Fabrick & Tabashnik, 2012; Li et al., 2019; Mathew et al., 2018; Morin et al., 2003; Wang et al., 2018; Wang et al., 2020; Wang et al., 2019b) Having available advanced tools, like this genome assembly, will facilitate the discovery and understanding of the underlying factors driving the evolution of resistance to Bt crops.

## Acknowledgements

This work was supported by the U.S. Department of Agriculture, Agricultural Research Service (USDA-ARS). The genome assembly was generated as part of the USDA-ARS Ag100Pest Initiative. Biological materials were provided by the USDA-ARS Pest Management & Biological Control Research Unit project number 2020-22620-023-000-D. This research used resources provided by the SCINet project of the USDA-ARS project number 0500-00093-001-00-D. JC and CH were supported, in part, by an appointment to the Research Participation Program at the Agricultural Research Service, United States Department of Agriculture, administered by the Oak Ridge Institute for Science and Education (ORISE) through an interagency agreement between the U.S. Department of Energy and USDA-ARS (contract numbers DE-AC05-06OR23100 and DOE-ARS-RPP-2019). The authors thank the members of the USDA-ARS Ag100Pest Team for sequencing and analysis support. All opinions expressed in this paper are the authors’ and do not necessarily reflect the policies and views of USDA. Mention of trade names or commercial products in this publication is solely for the purpose of providing specific information and does not imply recommendation or endorsement by the U.S. Department of Agriculture. USDA is an equal opportunity provider and employer.

## Data Availability

All raw reads used to assemble and scaffold this *Pectinophora gossypiella* genome were deposited at DDBJ/ENA/GenBank within BioProject PRJNA804955. These include the Sequence Read Archive (SRA) accession SRR17965739, which contains CLR data used to generate primary and alternate haplotig assemblies, and short paired end reads in accessions SRR18576017 and SRR21793435, used for contig polishing and Hi-C scaffolding, respectively. The primary ilPecGoss1.1 and ilPecGoss1.1 alternate haplotype assembly versions discussed in this manuscript are available in GenBank accessions GCF_024362695.1 (BioProject PRJNA804955) and GCA_024362825.1 (BioProject PRJNA836959), respectively. Both are under the Ag100Pest umbrella project, BioProject PRJNA555319. The primary assembly and annotations are also available at the i5k Workspace@NAL (Poelchau et al. 2015).

